# Methodology for isolation of serum, cerebrospinal fluid, and hippocampal neuron proteins from rat and their analysis using mass spectrometry-based shotgun proteomics

**DOI:** 10.1101/2024.05.15.594456

**Authors:** Pratibha Sharma, Rajinder K Dhamija

**Affiliations:** Human Behavior Department, Institute of Human Behaviour and Allied Sciences, New Delhi, India

## Abstract

Emerging interests in the field of research related to diseases such as Alzheimer’s disease, Parkinson’s disease, cognition, and other mental health-related disorders have prompted a need for a common method for the isolation of serum, CSF, and hippocampus. The hippocampus is responsible for learning and memory. It can be affected by various neurological and psychiatric disorders. However, the process of collecting samples such as CSF and hippocampal neurons is challenging, especially for small animals like rats. We have presented here a method for the isolation of serum, CSF, and hippocampal neurons that can be used for its downstream applications such as proteomics. We have used high-speed centrifugation instruments and density gradient centrifugation methods, which are easy to follow. Additionally, we have tested the proteins identified through mass spectrometry. Our method enables the study of proteins in serum, CSF, and neural cells for researching protein cross-talks and neurological disorder mechanisms.

## Introduction

Disruption of normal CSF (cerebrospinal fluid) contributes to the development of many diseases such as neurodegenerative disorders of Alzheimer’s disease, Parkinson’s disease, ischemic and traumatic brain injury, and neuroinflammatory conditions of multiple sclerosis [1]. CSF has shown potential usefulness in the diagnosis of Alzheimer’s and Parkinson’s disease from Abeta 1-42 and alpha-synuclein as diagnostic biomarkers respectively [2, 3]. During diseased conditions, neuro-inflammation is mediated by inflammatory cytokines [4-6]. The cytokines are circulated across the blood-brain barrier by a saturated transport mechanism. There is overwhelming evidence in support that the system’s immune response cross-talk with brain pathology [7]. CSF is the clear, colorless body fluid produced in the choroid plexus of the brain’s ventricles. It acts as a cushion in providing mechanical and immunological protection to the brain and spinal cord from injury. It is an important sample for diagnosing diseases in the central nervous system (CNS). However, injecting small animals and collecting CSF can be challenging and often results in high mortality rates.

The hippocampus plays an important role in learning and memory. The neurons get degenerate in this region are associated with neurodegenerative disorders and cognitive disorders [8, 9]. In Alzheimer’s disease, the neurons lose connections and neuronal damage occurs in the parts of the brain involved in memory, including the entorhinal cortex and the hippocampus. Shrinkage is severe in the hippocampus, an area of the cortex that plays a key role in the formation of new memories. The role of neural regeneration therapies has been implicated in these diseases [10]. Rodents are used majorly for laboratory work and sample collections because their genetic, biological, and behavioral characteristics closely resemble those of humans [11]. Also, they are cheap and breed quickly. The study of neurons is important for understanding the nervous system and neurodegeneration mechanism. It is challenging to obtain CSF from rats in appropriate amounts and without contamination by blood. However, this fluid is essential to study the protein interactions and mechanisms involved in various diseases, especially when combined with hippocampal neuron isolation. We have optimized the protein isolation and analysis utilizing mass spectrometry-based shotgun proteomics after simultaneous collection of serum, CSF, and hippocampal neurons from pre-anesthetized rats. Our method has a 90% success rate in producing clear CSF, as an improvement over the previous method [12, 13]. After serum and CSF collection, we immediately sacrificed rats to isolate the hippocampus and obtain pure neuron fractions. We used high-speed centrifugation with density gradients to isolate plasma membrane proteins from intracellular, mitochondria, and microsomal materials. Researchers may choose to work on these proteins depending on their research interests. Further, if required neuron culture can be performed [14]. We have meticulously devised a comprehensive sampling protocol that is specifically tailored to cater to various downstream applications such as proteomics. The methodology is designed to enable protein identification through mass spectrometry by sampling serum, CSF, hippocampal neuron intracellular, and plasma membrane from rats. With careful consideration of the various factors involved, we have ensured that the method is accurate, efficient, and reproducible.

## Materials and Methods

All procedures involving animals were approved by the Institute Ethical Committee of All India Institute of Medical Sciences, New Delhi (IAEC No.-647/IAEC/11).

### 1. Blood collection and serum isolation from Wister rat

1. A Wister albino strain of a male rat, weighing 180 g was used. The blood sample was collected with the help of an expert from the retrobulbar plexus or sinus of a rat eye without anesthesia. This procedure can be performed under general anesthesia as described previously [15, 16].
2. Hold the rat with the tail side and rotate the facing head towards gravity in a circular movement to increase the flow of blood. The rat was scuffed with the thumb and forefinger of the left hand and the skin around the eye was pulled tight.
3. A fresh and sterile capillary was inserted into the medial canthus of the eye at a 30-degree angle to the nose. Applied slight thumb pressure to puncture the tissue and enter the plexus or sinus.
4. Blood started coming to the capillary and was allowed to fall in clean and sterile 1.5 mL Eppendorf. After a few minutes, the same experiment may be repeated to collect blood from another eye. We collected approximately 1 ml of blood from one eye at a time.
5. For isolation, serum samples were kept undisturbed at 4oC and incubated for 1 h to allow the blood to clot. Blood samples were then subjected to centrifuge to remove the clot at 1000 g for 10 mins at 4°C.
6. 120 μL of clear serum was collected. Any blood-contaminated samples were discarded. Aliquots of 100 μL each of serum samples were collected and stored at - 80°C for further use.

### 2. CSF collection

1) Intraperitoneally anesthetized the above rat by injecting ketamine (100 mg/kg) and xylazine (10 mg/kg). Add 1X PBS, pH 7.4 up to 1 mL. Rat will not respond to toe pinch on getting anesthetized. Note: Prepare the dose according to the weight of the rat. PBS is added to prevent dehydration.
2) Specially constructed ear bars were placed in the external auditory meatus and the rat was fixed properly to the Stereotaxic instrument.
4) Flex the head downward at 90 degrees, and a depressible surface with the appearance of a rhomb between occipital protuberances and the spine of the atlas becomes palpable. Cleaned the head hair using scissors and a razor. The cleared head was wiped with 70% Alcohol.
5) At the midline of the scalp incision was made. The cervico-spinal muscle was reflected and the atlanto-occipital membrane was exposed.
6) The rat head was laid down at a 135-degree angle to the body. The atlanto-occipital membrane was punctured by the fire-polished 1ml syringe with a 27G needle. By gentle aspiration, non-contaminated CSF was collected in the syringe. CSF contaminated with blood was discarded. Note: We were able to collect 80-160 μL of CSF. The volume of CSF collected will vary depending on rat gender, age, size, and other health conditions.
7) CSF was centrifuged at 14,000 g for 15 min at 4°C to remove any cells present in it. Collected CSF incubated at 4°C for 1 h and stored in 50 μL aliquot each at -80°C for further use.

### 3.

Isolation of the hippocampus from the brain
1. After collecting the CSF, the rat was immediately sacrificed. Two mL of chloroform-soaked cotton were placed inside a container with the rat and closed tightly.
2. After approximately 20 seconds, the rat stopped the movement, removed by confirming anesthesia for a lack of withdrawal reflex from a toe pinch. Note: The above anaesthetization is generally effective for 25-30 min.
3. Decapitate with guillotine and disinfect the head with 70% ethanol. Dissected the skin over the top of the skull to expose it. Inserted scissors into the spinal canal and carefully cut the calvarium on one side nearly to the front. Similarly, repeat on the other side. Care was taken not to damage the brain.
4. With the help of forceps, the base of the skull was grabbed and carefully taken hippocampi. To dissect the hippocampi, we orient the brain at the dorsal side so that the clear midline of the two hemispheres is visible. With curved forceps, we inserted down the dorsal midline to approximately half the depth and squeezed to sever the cerebral commissure. Re-inserted with 2 mm spread into the midline of the midbrain. Gently peeled one hemisphere to the side. Repeated the above steps with the contra-lateral hippocampus.
5. Both the hippocampi from the rat brain were collected into 2 mL PBS at 4°C in a 35mm diameter petri dish. Washed three times with PBS.

Note: Care must be taken not to squeeze the tissue as this will affect the neuronal isolation.

### 4. Isolation of neurons

1. After removing the hippocampus, we placed it on a sterile filter paper that was pre-wet with PBS. We sliced the hippocampus into pieces of 0.5 mm. Then, we immediately transferred the tissue slices into a 15 mL corning tube containing 5 mL PBS, which was kept at 4°C.
2. Next, we placed the 15 mL corning tube with the tissue in a shaking water bath at 30°C for 10 minutes to ensure the temperature was consistent throughout.
3. For this we added 12 mg of papain powder to 6 mL of PBS in a 15 mL Corning tube. The tube was then kept in a 37°C shaking incubator for 30 min before use. Any insoluble particles were filtered out, and the resulting solution was sterilized and kept on ice until needed. This solution must be freshly prepared for every experiment.
4. Using a wide bore pipette, we transferred the tissue to a Corning tube containing papain at a temperature of 30°C. Shaked for 30 minutes at a rate of 170 rpm. Again, using a wide bore pipette, we transferred the tissue to a 15 mL Corning tube containing as little papain as possible in 2 mL of PBS at a temperature of 30°C. Let it sit for 5 minutes at room temperature. After 1 minute, transfer the supernatant to another empty 15 mL Corning tube.
5. Using a 9-inch wide bore Pasteur pipette, we triturated the contents approximately 10 times in 45 seconds. Each trituration step involved sucking the tissue up into a pipette, without air bubbles, and immediately emptying the contents back into the tube, without air bubbles.
6. We re-suspend the sediment from the first tube in 2 mL PBS. Repeated the above trituration steps two more times, combining the supernatants from each trituration.
7. The OptiPrep solution of 60% iodixanol in water with density 1.32 g/mL was used. In 15 ml Corning layered 7%, 10%, 12%, and 20% solution of OptiPrep in PBS. Layers from higher density solution at the bottom to the top were prepared. Carefully applied the cell suspension to the top of the prepared gradient. The cell suspension was floated on top of the gradient.
8. Centrifuged the gradient at 3000 g in a swing bucket centrifuge for 15 min at RT.
9. We aspirated the top 6ml containing cellular debris. The top 1 mL of fraction 1 was enriched with oligodendrocytes was aspirated.
10. We collected fraction 3 (from the lower edge of the dense band to 0.5 mL from the bottom of the tube) was enriched for neurons. Fraction 2, the opaque band, contains cell fragments, neurons, and other cells and fraction 4 (bottom 0.5ml and loose pellet) contains microglia.
11. Fraction 3 contained pure neurons, and was collected for further applications. To ensure the purity of the neurons, the step mentioned above is repeated. The fraction of pure neurons obtained was diluted in 5 mL PBS and centrifuged at 1500 g for 15 min.
12. The pellet obtained was washed thrice in PBS at 1000 rpm for 5 min each. Resuspended the pellet in 100 μL of resuspension buffer (0.25 M sucrose, 10 mM MgCl2, 5 mM Tris-pH 7.4, 1 mM PMSF)
13. We sonicated for five cycles with 20s ON and 40s OFF each for 5 min. We homogenized the suspension with ten strokes in five minutes.
14. The sample was centrifuged at 14,000 g at 4°C for 15 min. The supernatant was stored at -80°C for further applications. The obtained supernatant had intracellular proteins from hippocampal neurons. The pellet was processed further for plasma membrane protein isolation.

### 5. Isolation of plasma membrane proteins

1. The pellet was diluted in PBS. We prepared sucrose gradient in ultra-clear tubes. The sucrose gradients were prepared in PBS, pH-7.4, and layered bottom to top with 60%, 35.5%, 25.5%, and 19% respectively. The samples can be placed either at the bottom or at the top of the layer carefully using a Pasteur pipette of 3 mL by the side wall of the tube.
2. Ultracentrifuged at 1,00,000 g for 1:45 hr at 4°C in Beckman Coulter SW Ti 41 rotor by overlaying onto sucrose density gradient. We obtained plasma membrane proteins at the interface of 25.5% and 35.5% of sucrose.
3. We collected the plasma membrane fraction and diluted it in PBS. We repeated the ultracentrifugation step at 100,000 g at 4°C for 1:45 hrs using a fixed angle rotor.
4. We discarded the supernatant and collected the pellet. The obtained pellet contained hippocampal neuron plasma membrane proteins. The pellet was stored at -80°C for further experiments.

### 6. Protein precipitation by acetone precipitation method

1. Four times the volume of cold (-20°C) acetone was added to the CHAPS-solubilized samples. We vortexed and incubated the tubes for 45 min at -20°C. It was centrifuged at 15,000 rpm, 4°C for 10 min.
2. The supernatant was discarded without disturbing the protein pellet. We re-suspended the protein pellet in 80% acetone (pre-chilled) and centrifuged as above.
3. The above steps were repeated twice to obtain the protein pellet. We dried off the acetone completely from the protein pellets.

### 7. Protein quantification

1. The protein pellet was dissolved in 1 mL of 10 mM PB, pH 7.4, 0.1 % Triton-X-100. A protease inhibitor cocktail was added to all samples to avoid proteolysis according to sample volume.
2. Protein concentration in the protein sample was determined by using a BCA protein assay kit (Pierce, Thermo Scientific). Samples in aliquots of 100 μL each were stored at -80°C.

### 8. 2D gel electrophoresis

1. The samples of serum, CSF, and hippocampal tissue proteins from intracellular, and plasma membranes were collected at 200 μg each. These were further solubilized in lysis buffer (8M Urea, 3M Thiourea, 4% CHAPS). Plasma membrane samples were solubilized carefully in a solubilization buffer.
2. Membrane preparations were resuspended (0.5 mg/mL) in 20 mM HEPES, pH 7.4, 150 mM KCl, 1 mM EDTA, 1 % (w/v) CHAPS and solubilized at 4 ºC for 1 h with end-to-end rotation.
3. The solubilized membrane was centrifuged at 15,000 rpm for 15 min, and supernatants were collected.

#### 8.1 Sample rehydration

1. 1.25 μL of IPG buffer (pH 3-10 NL) (Amersham Biosciences, USA), 1μl of BPB (from 0.002% stock), and 0.75 mg of DTT were added to each sample.
2. We made a final volume of 250μl with lysis buffer. After mixing, the tube containing samples was centrifuged the samples at 15,000 rpm for 2 min and loaded on a rehydration tray (Amersham Biosciences, USA).
3. IPG-strip of pH range 3-10 NL, 13 cm was used for IEF. The strips were placed carefully over the sample for rehydration for 14-16 hrs.

#### 8.2 Two-Dimensional gel electrophoresis (Iso-electric focusing)

1. We rehydrated the IPG strip for iso-electric focusing in an IPGphor 3 (Amersham Biosciences, USA). We used total volt-hours of 30,000 VhT.
2. The strips were equilibrated in SDS-equilibration buffer containing 50 mM Tris-HCl, pH 8.8, 6 M urea, 30 % glycerol, 2 % SDS, and 0.02 % bromophenol blue.
3. The strips were first equilibrated for 15 min with 0.05 % DTT prepared in 5 mL of SDS equilibration buffer at RT. The solution was decanted and replaced with 1.25 % iodoacetamide solution, prepared in 5ml of SDS equilibration buffer, for 15 min at RT.
4. 10% polyacrylamide gels were prepared using a Ruby gel apparatus (Amersham Biosciences, USA). The strips were carefully loaded on the PAGE and sealed with the help of an agarose solution (0.5% agarose, 0.002% BPB).
5. The gel was run at 15 mA for 30 min followed by 30 mA till the bromophenol blue tracking dye came out of the gel. The gels were first stained using Colloidal Coomassie. We destained the colloidal Coomassie and re-stained using silver stain for compatibility with mass spectrometry.

### 9. Protein digestion by In-solution trypsinization of proteins

1. We dissolved the sample pellet in 100 μL of 50mM Ammonium bicarbonate and adjusted pH 8.0. 5ul of DTT (prepared in 100 mM Ammonium bicarbonate) was added. We used 100 μg of each serum, CSF, hippocampal intracellular protein, and plasma membrane protein samples for further processing.
2. We boiled the samples for 10 min and kept them at RT for 1hr after gentle vortex and spin.
3. We added 4 μL of Iodoacetamide, vortexed, spun, and kept at RT for 1 hr.
4. We added 2 μL (1μg/μL) of trypsin on ice to each sample. Vortexed, spun, and incubated at 37°C for 16 h. We lyophilized the samples. The samples were lyophilized to dry up completely. Peptide desalting was performed before mass spectrometry analysis.

### 10. Peptide desalting, LC-MS/MS analysis, and data analysis

1) We purchased peptide desalting spin columns (Pierce, Thermo Scientific) and used them as per manufacturer protocol.
2) The dried peptides were resuspended in 0.1% Formic acid.
3) We used 2 μL of each sample peptide to quantify concentration with a Nanodrop instrument.
4) We used label-free quantitation and diluted 2 μg sample peptide in 10 μL of 0.1% FA.
5) We equilibrated the column and analytical column with 0.1% FA for LC parameters. The samples were kept in an autosampler. 1 μg of each digested peptide was injected into the column.
6) The LC gradient program was set according to the sample complexity using gradients of 125 min. MS parameters were optimized using standard peptides. After optimization, we run the program.
7) We obtained the raw data files of the MS/MS spectrum and analyzed them using Mascot software for protein analysis.
9) Further bioinformatics analysis of Gene ontology and pathway were performed.

## Results and discussions

### Isolation of serum and CSF proteins

We have optimized the method for the isolation of proteins from serum, CSF, and brain hippocampal neuron proteins for downstream applications such as proteomics. We have used thirty Wister albino rat strains. All of these were tested for serum, CSF, and hippocampal neuron isolation. Serum is the clear, colorless blood plasma not including the fibrinogens (it neither contains white or red blood cells nor the clotting factors). Multiple studies have identified serum and CSF proteins whose expression levels in Alzheimer’s, Parkinson’s disease, and dementia patients differ from controls [17-20]. The advantages of blood serum for biomarker investigations are its ease of sampling and collection, as well as its minimal contamination. Using the above method, it is possible to collect 1-2 mL of blood from a rat per session. Clear CSF can also be collected without any contamination using general anesthesia. Ketamine, a medication that provides amnesia, analgesia, and dissociation from the environment, is often given with xylazine, an α2 adrenergic agonist. The combination of ketamine and xylazine provides relatively safe, short-term anesthesia [12]. To simplify the method for maximum collection of CSF from cisterna magna, the rat was fixed properly to a stereotaxic instrument shown in Figure 1. We successfully aspirated 3.1 mL (varied from 80-160 μL per animal of weight range 180-240 g including male and female, 15 each). The success rate was 90% by the above method except for two CSF samples that were blood contaminated and one faced needle obstruction. These rats after collection of CSF samples were sacrificed for isolation of hippocampus [Figure 1].

**Figure 1:**
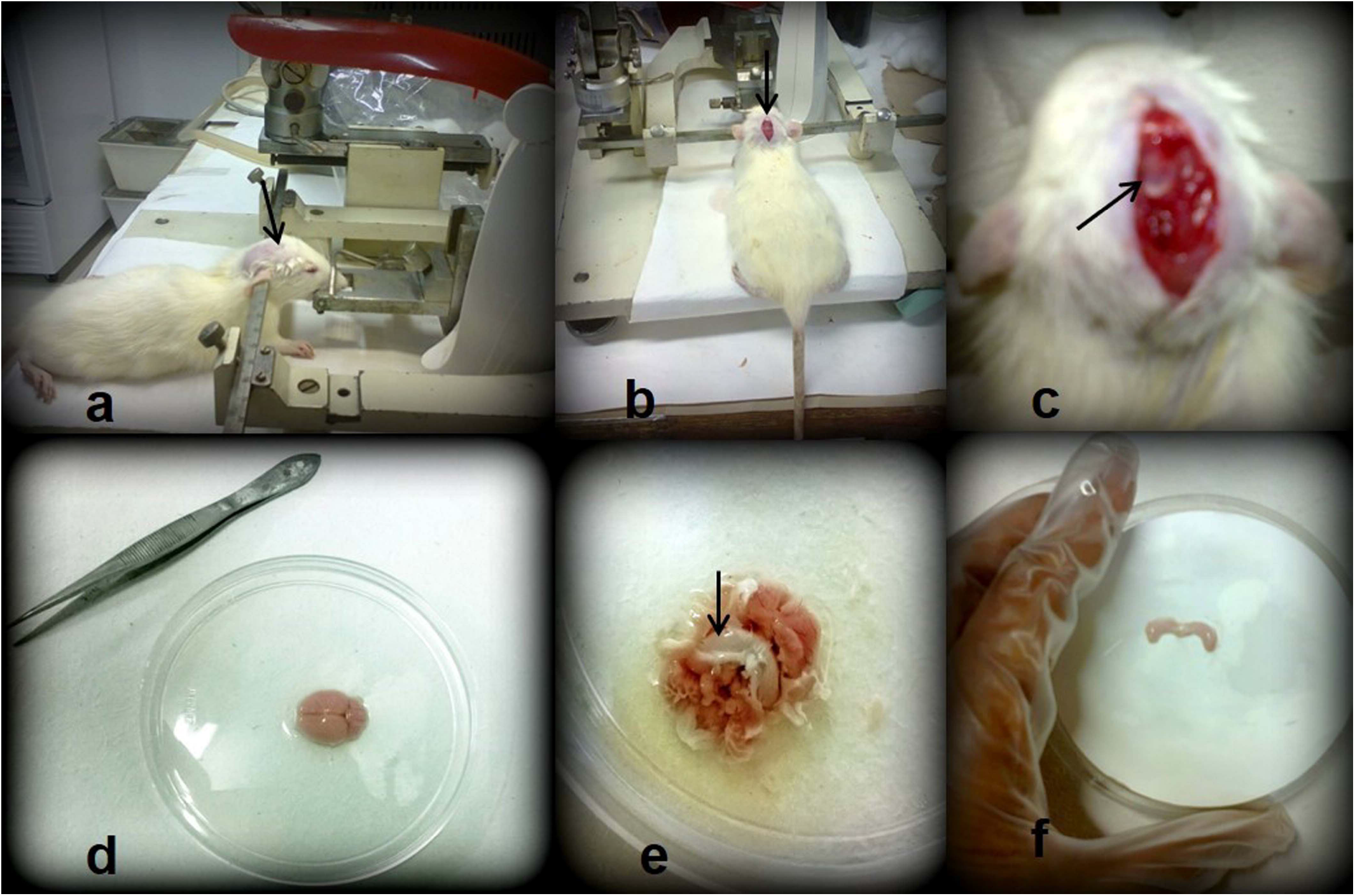
Methodological steps for collection of CSF and isolation of brain hippocampus. Collection of CSF (a,b,c) and isolation of hippocampus tissue (d,e,f). a. Wister albino rat was fixed in stereotaxic instrument for CSF collection. b. Cleared head hair and incision is made at the midline of scalp. Cervico-spinal muscle reflected. c. atlanto-occipital membrane is exposed., punctured for CSF collection. d. Isolated rat brain. e. hippocampus tissue is exposed, f. isolated hippocampus.

### Isolation of hippocampal neuron proteins

The optical density gradient method was used to separate cell debris and microglia from the neuron fraction [Figure 2]. Fraction 2 contains the hippocampal neurons, but it was of low purity and may be contaminated by other cells. Fraction 3 of the gradient contains pure neuron fraction obtained from one rat hippocampal tissue. The triturating step is very critical in neuron isolation. The hippocampal cut slices treated with papain were triturated carefully and gently. Obtained neuron cells were dissolved in a resuspension buffer and cells were lysed using sonication. Sonication works by applying sound energy to agitate particles in the sample disrupt the neuron cell membranes and release cytosolic components. It speeds up dissolution by breaking the intermolecular interactions. The cell suspension was homogenized. This helped in extracting membrane proteins and solubilizing protein complexes to preserve the protein-protein interactions. Subsequently, the membrane fraction was purified on a discontinuous sucrose gradient between 35% and 25.5% sucrose [Figure 2]. Mitochondrial and ER fractions were removed (at an interface of 35.5% and 50% gradient) from the plasma membrane. Sucrose are large carbohydrate molecules and their presence may interfere with protein-protein interaction studies. These were removed either by dialysis or protein precipitation methods. Re-ultracentrifugation was followed by diluting the plasma membrane fractions in PBS at pH-7.4, such that the final concentration of sucrose was less than two percent. 1mM PMSF was added to the membrane protein commonly used in protein-solubilization to deactivate proteases from digesting proteins of interest after cell lysis.

**Figure 2:**
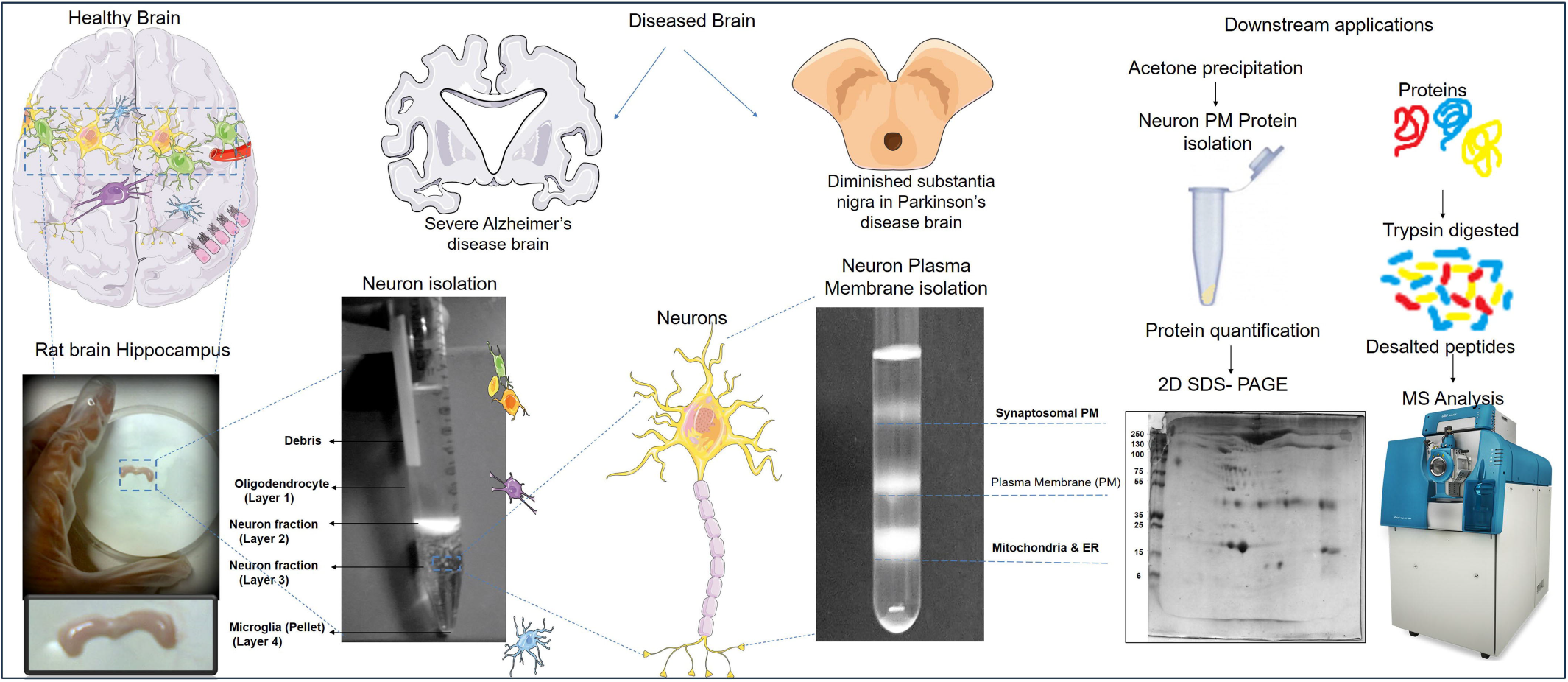
Isolation of hippocampus and neuron proteins for downstream applications. Isolation of rat brain hippocampal neuron-using optiprep density gradient. Fraction 3 contains pure neuron fraction, repeated twice with fraction 3 to obtain more purity of neurons. Isolation of Hippocampal neuron plasma membrane protein-sucrose density. Isolation of hippocampal neuron intracellular and plasma membrane proteins for mass spectrometry protein identification and other downstream applications. Cartoon images of brain and neurons were obtained from SMART-Servier Medical ART available at https://smart.servier.com/.

### Downstream application of isolated proteins

#### 2D-SDS PAGE and enzymatic digestion

Two-dimensional sodium dodecyl sulfate-polyacrylamide gel electrophoresis (2D SDS-PAGE) is a reliable and efficient method for the separation of several hundreds to a few thousands of proteins. The proteins were separated firstly based on differences in their isoelectric point (pI) and secondly based on molecular weight [Figure 3]. In-solution digestion in comparison to in-gel digestion provides more control over the outcome of the MS analysis in generating a greater number of peptides. Low molecular weight proteins are mostly difficult to identify from the gels. The recovery of peptides and alterations in conditions like pH, protein concentration, digestion buffer, and proteolytic enzyme is easier. However, this may result in the loss of proteins during processing and samples need to be salt-free before MS analysis. These methods can achieve femtomole quantities of proteins. Trypsin digested peptides were identified by mass spectrometry.

**Figure 3:**
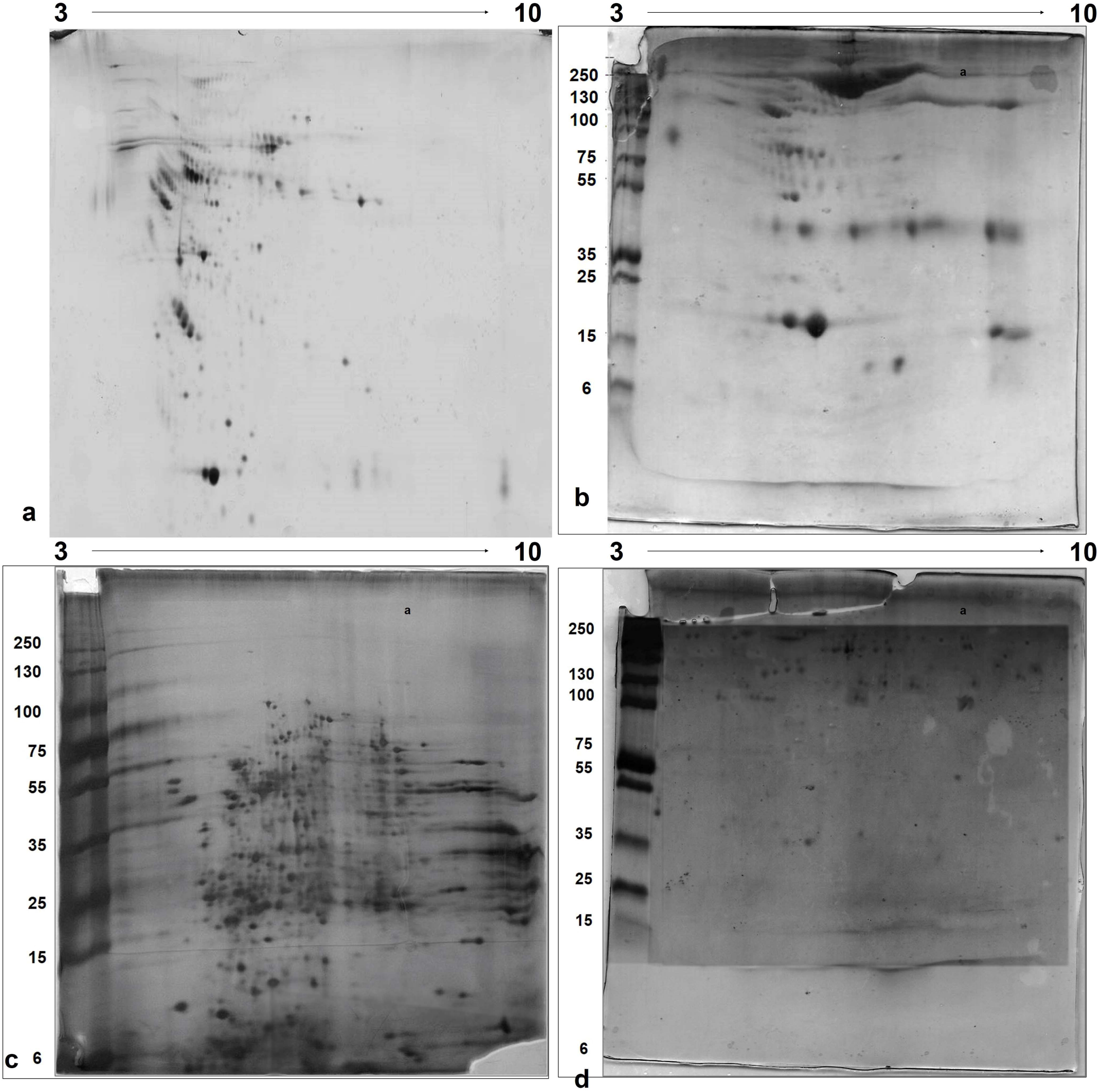
2D-PAGE of serum, cerebrospinal fluid, hippocampal neuron intracellular proteins, and hippocampal neuron plasma membrane proteins. 10% 2D-PAGE gel, pH 3-10 NL, 13cm strip. Stained with silver stain.

### Protein identification by mass spectrometry

We used ESI Triple TOF 5600 (SCIEX) to identify trypsin-digested peptides from serum, CSF, hippocampal neuron intracellular, and plasma membrane proteins. This tool is exceptionally useful for analyzing complex proteomics samples. The database of Rattus norvegicus species was searched using the software Protein Pilot vs. 4.2. We have identified 115, 92, 147, and 173 numbers of proteins in the serum, CSF, HNPM & HNIC respectively, with less than a 1% false discovery rate. A non-redundant list of these identified proteins has been listed in Supplementary Table S1-S4. The protein scores result from the MS/MS result search are derived from the ions scores.

### Protein-protein interaction analysis

We performed protein-protein interaction of proteins identified by mass spectrometry using Cytoscape vs. 3.9.1 and String plugin. We observed many proteins similar in the CSF, serum, and hippocampal intracellular proteins. Protein-protein interaction networks from four samples merged and individual maps have been presented in Figure 4.

**Figure 4:**
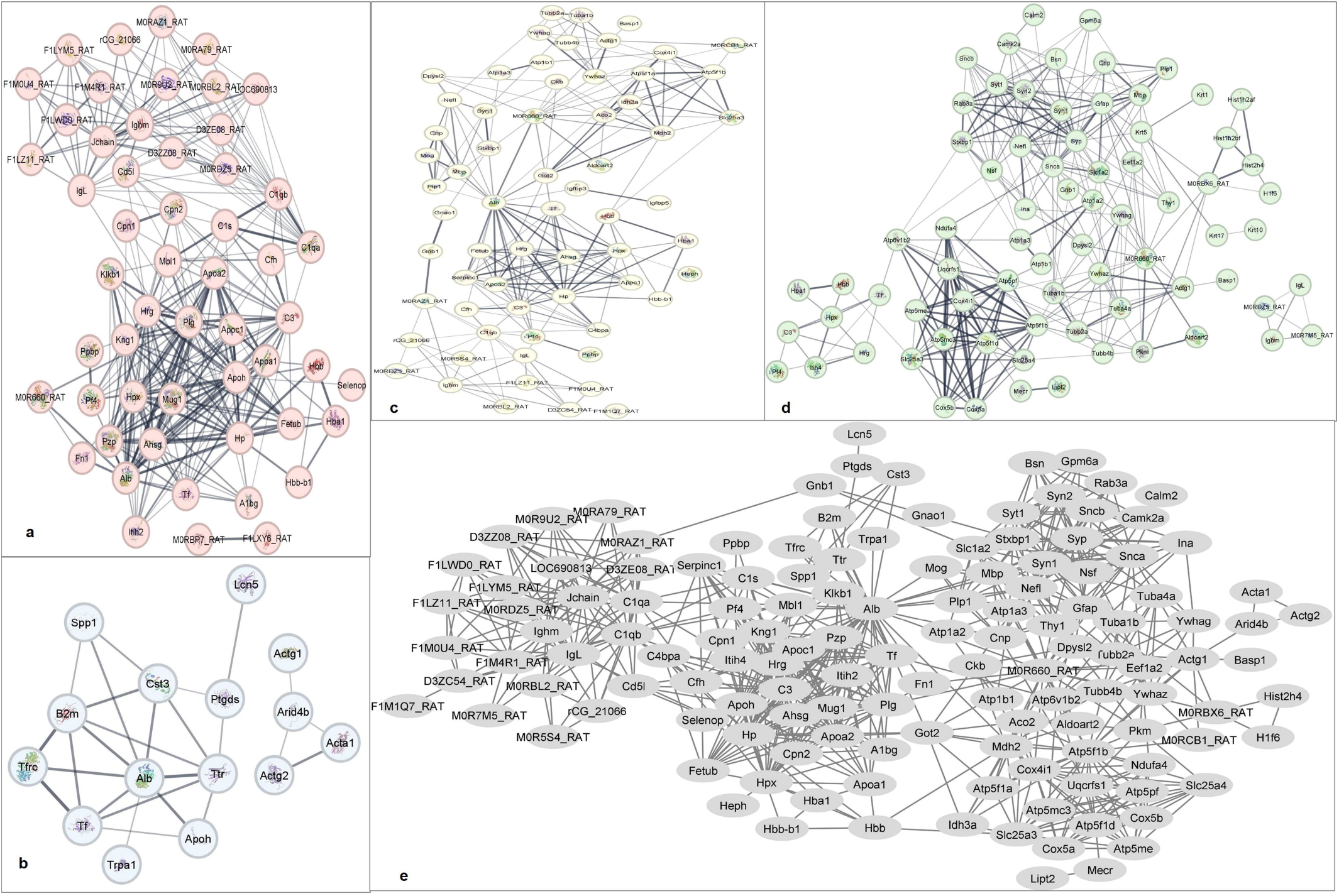
Protein-protein interactions study of serum, CSF, hippocampal neuron intracellular, and plasma membrane proteins identified by mass spectrometry. Serum proteins (a), CSF (b), hippocampal neuron intracellular (c), hippocampal neuron plasma membrane proteins (d). Protein nitration networks obtained from CSF, serum, hippocampal neuron intracellular and membrane proteins identified were merged to show protein-protein interactions (e). This interaction data was obtained using Cytoscape software and String.

### Implication in disease mechanism

Transferrin and albumin were found in both CSF and serum and their dysfunction might have an association with neurodegenerative disorders [21-24]. CSF is in direct contact with the extracellular space of the brain and can reflect biochemical changes that occur inside the brain [4-6]. CSF and serum have 80% common proteins. For these, reasons both the CSF and serum are considered for the biomarker discoveries which can help in early diagnosis of Alzheimer’s disease. Blood-brain barrier (BBB) dysfunction allows the entry of blood proteins into CSF. The CSF flow abnormality can result in impaired absorption of proteins and elevations of acute inflammatory mediators, including cytokines reported in neurodegenerative disorders. Cytokines are circulated across the BBB by a saturated transport mechanism and cross-talk with brain pathology [7]. In Alzheimer’s disease, the neurons lose connections and neuron damage occurs in the parts of the brain involved in memory, including the entorhinal cortex and the hippocampus [8, 9]. Shrinkage is severe in the hippocampus, an area of the cortex that plays a key role in the formation of new memories. This method provides the best approach to studying plasma membranes, glial cell-free, and rich fractions from hippocampal neurons of the central nervous system. We have developed a combined method for the collection of CSF and hippocampus neurons with an improved success rate in producing clear CSF. The entire process of CSF isolation and hippocampus neuron isolation will take approximately four hours. We have isolated pure neuron fractions by repeated ultracentrifugation and density gradient centrifugation steps. We have successfully isolated neural cell plasma proteins and identified them using mass spectrometry. Our method is cost-effective, simple, and reproducible. Our method enables the study of proteins in serum, CSF, and neural cells for researching protein cross-talks and neurological disorder mechanisms.

## Acknowledgments

We would sincerely like to acknowledge Prof. Alagiri Srinivasan, Professor (Full), Department of Biophysics, All India Institute of Medical Sciences, New Delhi for his supervision and intellectual discussions. This project work was supported by a fellowship grant from the Indian Council of Medical Research Code-13340.

## Notes

### Competing Interest Statement

The authors have declared no competing interest.

