## Supplementary material for "Methodology for isolation of serum, cerebrospinal fluid, and hippocampal neuron proteins from rat and their analysis using mass spectrometry-based shotgun proteomics": Supplementary data.docx

**Table S1:** Proteins identified by mass spectrometry from serum sample

| **Total**  **Score** | **% Cov (95)** | **Uniprot id** | **Protein Name** | **Peptides (95%)** |
| --- | --- | --- | --- | --- |
| 118.49 | 36.7 | M0RBJ7 | Complement C3 | 63 |
| 61.48 | 40.4 | P12346 | Serotransferrin | 33 |
| 48.80 | 50.7 | P20059 | Hemopexin | 30 |
| 60.02 | 14.8 | P04937 | Fibronectin | 29 |
| 36.44 | 49.5 | Q5M7V3 | LOC367586 protein | 29 |
| 33.23 | 41.9 | Q5M842 | IgG-2a protein | 23 |
| 39.86 | 17.7 | Q71SA3 | Thrombospondin 1 | 19 |
| 18.69 | 24.8 | Q4VBH1 | Ighg protein | 18 |
| 18.69 | 26.5 | Q569B4 | Ighg protein | 18 |
| 29.35 | 25.8 | P08934 | Kininogen-1 | 16 |
| 27.82 | 24.8 | Q569B3 | Igh-6 protein | 16 |
| 29.89 | 27.6 | P02770 | Serum albumin | 15 |
| 16.89 | 27.8 | Q4KM66 | LOC500183 protein | 14 |
| 25.62 | 31.2 | Q5M8A0 | Kng2 protein | 14 |
| 25.14 | 37.5 | P06866 | Haptoglobin | 13 |
| 10.75 | 63.2 | P01836 | Ig kappa chain C region | 11 |
| 20.90 | 5.5 | Q6MG79 | Complement component 4 | 9 |
| 10.98 | 56.3 | B1H216 | Hemoglobin alpha | 7 |
| 12.24 | 5.4 | G3V9R2 | Protein Cfh | 7 |
| 12.20 | 42.2 | P02091 | Hemoglobin subunit beta-1 | 7 |
| 12.01 | 14.1 | Q99PS8 | Histidine-rich glycoprotein | 7 |
| 9.41 | 33.3 | P11517 | Hemoglobin subunit beta-2 | 6 |
| 9.65 | 53.9 | P20767 | Ig lambda-2 chain C region | 6 |
| 8.97 | 42.1 | P19939 | Apolipoprotein C-I | 5 |
| 9.75 | 3.9 | Q63041 | Alpha-1-macroglobulin | 5 |
| 6.14 | 5.1 | D3ZFH5 | Protein Itih2 | 4 |
| 7.21 | 41.2 | F1M0U4 | RCG21092 | 4 |
| 8.18 | 11.9 | Q5U3X5 | Fgl2 protein | 4 |
| 7.13 | 10.1 | Q6P7S6 | Clusterin | 4 |
| 6.73 | 12.8 | Q6PAH0 | Apolipoprotein E | 4 |
| 5.14 | 26.6 | F1LZ11 | Uncharacterized protein | 3 |
| 2.29 | 23.1 | F8SQR6 | Immunglobulin heavy chain variable region | 3 |
| 6.00 | 33.3 | P04638 | Apolipoprotein A-II | 3 |
| 5.85 | 37.1 | P06765 | Platelet factor 4 | 3 |
| 4.73 | 11.7 | P24090 | Alpha-2-HS-glycoprotein | 3 |
| 6.55 | 9.0 | Q4KM75 | CD5 antigen-like | 3 |
| 4.87 | 11.6 | Q64599 | Hemiferrin | 3 |
| 4.70 | 18.9 | Q6PDV1 | Lysozyme 2 | 3 |
| 4.17 | 5.5 | Q9EPH1 | Alpha-1B-glycoprotein | 3 |
| 4.63 | 7.0 | Q9EQV8 | Carboxypeptidase N catalytic chain | 3 |
| 4.48 | 17.1 | E9PSU8 | Uncharacterized protein | 2 |
| 3.12 | 0.6 | F1LMV6 | Protein Dsp | 2 |
| 2.86 | 17.7 | F1M1R0 | Uncharacterized protein | 2 |
| 2.88 | 11.4 | F1M229 | Uncharacterized protein | 2 |
| 2.35 | 14.6 | F1M3Y4 | Protein RGD1564184 | 2 |
| 3.11 | 14.0 | G3V8Z5 | Uncharacterized protein | 2 |
| 2.23 | 19.3 | M0R628 | Uncharacterized protein | 2 |
| 1.75 | 0.7 | M0R6Z9 | Protein Myo18b | 2 |
| 4.17 | 7.7 | P04639 | Apolipoprotein A-I | 2 |
| 4.57 | 8.8 | P19999 | Mannose-binding protein A | 2 |
| 3.83 | 4.7 | P25236 | Selenoprotein P | 2 |
| 1.80 | 1.3 | Q03626 | Murinoglobulin-1 | 2 |
| 3.22 | 3.3 | Q5FVS2 | Kallikrein B | 2 |
| 3.21 | 4.3 | Q6QI47 | LRRGT00161 | 2 |
| 4.00 | 2.0 | Q7TMA9 | Aa1249 | 2 |
| 1.61 | 1.2 | Q91YB6 | Complement inhibitory factor H | 2 |
| 1.73 | 19.8 | Q99ME0 | CXC chemokine RTCK1 | 2 |
| 2.11 | 2.7 | B0BMT0 | RCG47746, isoform CRA_a | 1 |
| 0.72 | 4.0 | D3Z8A1 | REVERSED Protein Tctex1d1 | 1 |
| 0.48 | 5.3 | D3ZE08 | Uncharacterized protein | 1 |
| 2.00 | 11.1 | D3ZEP5 | Protein Igkv19-93 | 1 |
| 0.68 | 0.6 | D3ZH42 | REVERSED Protein Mov10l1 | 1 |
| 0.90 | 6.5 | D3ZPL2 | Uncharacterized protein | 1 |
| 0.57 | 1.2 | D3ZWU1 | Bromodomain containing 3 | 1 |
| 0.52 | 0.9 | D4AC99 | CTF18 | 1 |
| 2.00 | 2.9 | F1LPR6 | Uncharacterized protein | 1 |
| 3.36 | 2.3 | F1LQT4 | Protein Cpn2 | 1 |
| 2.00 | 13.0 | F1LWD0 | Uncharacterized protein | 1 |
| 2.04 | 18.5 | F1LYM5 | Uncharacterized protein | 1 |
| 2.00 | 10.3 | F1LYU4 | Uncharacterized protein | 1 |
| 1.42 | 9.6 | F1M4R1 | Uncharacterized protein | 1 |
| 2.11 | 7.7 | F1M5L5 | Uncharacterized protein | 1 |
| 1.72 | 10.4 | F1M663 | Uncharacterized protein | 1 |
| 2.00 | 2.8 | F1M6N0 | Protein LOC686143 | 1 |
| 2.57 | 9.4 | G3V6G1 | Immunoglobulin joining chain | 1 |
| 1.62 | 1.7 | G3V7L3 | Complement C1s subcomponent | 1 |
| 2.00 | 4.7 | G3V7N9 | Complement C1q subcomponent subunit B | 1 |
| 2.56 | 0.9 | G3V9J1 | Uncharacterized protein | 1 |
| 0.62 | 2.4 | M0R660 | Protein RGD1565368 | 1 |
| 2.00 | 5.7 | P31720 | Complement C1q subcomponent subunit A | 1 |
| 2.00 | 8.5 | Q32PY8 | Protein Sbsn | 1 |
| 2.00 | 2.5 | Q4G030 | C1r protein | 1 |
| 1.46 | 1.4 | Q4V8G6 | Methyltransferase-like 3 | 1 |
| 0.97 | 2.9 | Q5I0M1 | Apolipoprotein H | 1 |
| 0.67 | 11.8 | Q5RK13 | Igf1 protein | 1 |
| 1.44 | 2.3 | Q6IRS6 | Fetub protein | 1 |
| 1.80 | 1.2 | Q7TP84 | Ab1-346 | 1 |
| 1.65 | 0.5 | Q7TPK2 | Ac2-120 | 1 |
| 0.61 | 5.8 | Q9Z1I0 | Shc transforming protein | 1 |

**Table S2:** Proteins identified by mass spectrometry from CSF sample

| **Total**  **Score** | **% Cov (95)** | **Uniprot id** | **Protein Name** | **Peptides (95%)** |
| --- | --- | --- | --- | --- |
| 0.09 | 1.0 | D3ZNR4 | Sushi, nidogen and EGF-like domain-containing protein1 | 36 |
| 16.06 | 22.3 | Q8NHM4 | Putative trypsin-6 | 17 |
| 15.29 | 17.4 | A1A508 | PRSS3 protein | 14 |
| 11.64 | 18.8 | A8CED3 | Trypsinogen 5 | 12 |
| 11.64 | 18.8 | F8W7P3 | Trypsin-3 | 12 |
| 11.64 | 17.3 | P35030-2 | Isoform B of Trypsin-3 | 12 |
| 11.64 | 18.2 | P35030-3 | Isoform C of Trypsin-3 | 12 |
| 11.64 | 17.2 | P35030-4 | Isoform D of Trypsin-3 | 12 |
| 11.64 | 18.2 | Q6ISJ4 | Mesotrypsinogen | 12 |
| 11.64 | 17.2 | Q7Z5F4 | Protease serine 4 isoform B | 12 |
| 11.64 | 17.9 | Q8N2U3 | PRSS3 protein | 12 |
| 15.62 | 13.9 | P13645 | Keratin, type I cytoskeletal 10 | 8 |
| 3.85 | 10.3 | P22057 | Prostaglandin-H2 D-isomerase | 6 |
| 11.01 | 8.8 | P35908 | Keratin, type II cytoskeletal 2 epidermal | 6 |
| 10.00 | 20.2 | Q5NV56 | Anionic trypsinogen | 6 |
| 6.02 | 52.1 | P81605-2 | Isoform 2 of Dermcidin | 5 |
| 4.00 | 10.2 | Q6GMX3 | IGL@ protein | 2 |
| 2.00 | 6.4 | A0A5E4 | Uncharacterized protein | 1 |
| 2.00 | 14.3 | A0M8Q9 | C1 segment protein | 1 |
| 1.13 | 38.1 | A0N4V7 | HCG2039797 | 1 |
| 2.59 | 6.4 | A2NUT2 | Lambda-chain (AA -20 to 215) | 1 |
| 2.00 | 4.1 | A6XMV6 | Secreted phosphoprotein 1 | 1 |
| 0.15 | 0.0 | B4DZK1 | cDNA FLJ50036, weakly similar to Myosin-11 | 1 |
| 2.00 | 4.0 | B7Z351 | cDNA FLJ54682, highly similar to Osteopontin | 1 |
| 2.00 | 7.0 | B9A064 | Immunoglobulin lambda-like polypeptide 5 | 1 |
| 2.00 | 4.8 | C4B6Q2 | Osteopontin | 1 |
| 2.59 | 6.5 | C6KXN3 | Lambda light chain of human immunoglobulin surface antigen-related protein | 1 |
| 2.00 | 4.5 | F2YQ21 | Osteopontin-D | 1 |
| 0.15 | 0.9 | F5H149 | Coiled-coil domain-containing protein 180 | 1 |
| 6.02 | 12.5 | G3V7Q8 | Cationic trypsinogen | 1 |
| 10.00 | 20.2 | [P00763](http://www.uniprot.org/uniprot/P00763) | Anionic trypsin-2 | 1 |
| 2.00 | 16.0 | P01834 | Ig kappa chain C region | 1 |
| 2.00 | 8.8 | P02767 | Transthyretin | 1 |
| 65.34 | 48.8 | P02770 | Serum albumin | 1 |
| 2.00 | 8.8 | P06911 | Epididymal-specific lipocalin-5 | 1 |
| 2.31 | 8.4 | P07151 | Beta-2-microglobulin | 1 |
| 2.59 | 6.4 | P08721 | Osteopontin | 1 |
| 2.00 | 14.1 | P0CF74 | Ig lambda-6 chain C region | 1 |
| 2.00 | 14.1 | P0CG04 | Ig lambda-1 chain C regions | 1 |
| 2.00 | 14.1 | P0CG05 | Ig lambda-2 chain C regions | 1 |
| 2.00 | 14.1 | P0CG06 | Ig lambda-3 chain C regions | 1 |
| 2.00 | 4.1 | P10451 | Osteopontin | 1 |
| 2.00 | 4.3 | P10451-2 | Isoform B of Osteopontin | 1 |
| 2.00 | 4.5 | P10451-3 | Isoform C of Osteopontin | 1 |
| 2.00 | 4.5 | P10451-4 | Isoform D of Osteopontin | 1 |
| 2.00 | 4.3 | P10451-5 | Isoform 5 of Osteopontin | 1 |
| 9.00 | 9.5 | P12346 | Serotransferrin | 1 |
| 6.04 | 19.2 | P14841 | Cystatin-C | 1 |
| 1.82 | 2.6 | P26644 | Beta-2-glycoprotein 1 | 1 |
| 2.00 | 4.2 | P35572 | Insulin-like growth factor-binding protein 6 | 1 |
| 1.43 | 5.1 | P63259 | Actin, cytoplasmic 2 | 1 |
| 1.43 | 2.7 | P63269 | Actin, gamma-enteric smooth muscle | 1 |
| 1.43 | 3.9 | P68136 | Actin, alpha skeletal muscle | 1 |
| 2.00 | 7.8 | Q0KKI6 | Immunoblobulin light chain | 1 |
| 2.00 | 4.8 | Q3LGB0 | Osteopontin | 1 |
| 2.00 | 6.4 | Q567P1 | IGL@ protein | 1 |
| 2.00 | 6.4 | Q5CZ94 | Putative uncharacterized protein DKFZp781M0386 | 1 |
| 2.00 | 7.3 | Q5EFE6 | Anti-RhD monoclonal T125 kappa light chain | 1 |
| 2.00 | 6.5 | Q5FWF9 | IGL@ protein | 1 |
| 2.00 | 6.3 | Q6DHW4 | Uncharacterized protein | 1 |
| 2.00 | 6.4 | Q6GMV7 | Uncharacterized protein | 1 |
| 2.00 | 6.4 | Q6GMV8 | Uncharacterized protein | 1 |
| 2.59 | 6.4 | Q6GMW3 | IGL@ protein | 1 |
| 2.00 | 6.4 | Q6GMW4 | IGL@ protein | 1 |
| 2.00 | 6.4 | Q6GMW6 | Uncharacterized protein | 1 |
| 2.00 | 7.2 | Q6GMX0 | Uncharacterized protein | 1 |
| 2.59 | 6.4 | Q6GMX4 | IGL@ protein | 1 |
| 2.00 | 6.4 | Q6IN99 | IGL@ protein | 1 |
| 2.59 | 6.4 | Q6IPQ0 | IGL@ protein | 1 |
| 2.00 | 6.4 | Q6NS95 | IGL@ protein | 1 |
| 2.00 | 6.4 | Q6P2J1 | Uncharacterized protein | 1 |
| 2.00 | 6.4 | Q6P5S3 | Uncharacterized protein | 1 |
| 2.00 | 7.2 | Q6P5S8 | IGK@ protein | 1 |
| 2.00 | 6.4 | Q6PIK1 | IGL@ protein | 1 |
| 2.00 | 7.2 | Q6PIL8 | IGK@ protein | 1 |
| 2.59 | 6.4 | Q6PIQ7 | IGL@ protein | 1 |
| 2.00 | 7.2 | Q6PJF2 | IGK@ protein | 1 |
| 2.59 | 6.4 | Q6PJG0 | Uncharacterized protein | 1 |
| 2.00 | 10.8 | Q6PJR7 | IGL@ protein | 1 |
| 0.15 | 1.2 | Q6QD51 | Coiled-coil domain-containing protein 80 | 1 |
| 1.43 | 0.9 | Q6RI86 | Transient receptor potential cation channel subfamily A member 1 | 1 |
| 2.00 | 6.4 | Q7Z2U7 | Uncharacterized protein | 1 |
| 2.59 | 6.4 | Q8N355 | IGL@ protein | 1 |
| 2.59 | 6.4 | Q8N5F4 | IGL@ protein | 1 |
| 2.00 | 6.4 | Q8NEJ1 | Uncharacterized protein | 1 |
| 2.00 | 7.1 | Q8TCD0 | Uncharacterized protein | 1 |
| 2.00 | 14.1 | Q8TCJ5 | Putative uncharacterized protein DKFZp667J0810 | 1 |
| 0.09 | 0.5 | Q8TER0-4 | Isoform 3 of Sushi, nidogen and EGF-like domain-containing protein 1 | 1 |
| 2.00 | 6.4 | Q96E61 | Uncharacterized protein | 1 |
| 9.00 | 9.5 | Q99376 | Transferrin receptor protein 1 | 1 |
| 0.15 | 0.9 | Q9JKB5 | AT-rich interactive domain-containing protein 4B | 1 |
| 0.15 | 1.2 | Q9P1Z9-4 | Isoform 4 of Coiled-coil domain-containing protein 180 | 1 |

**Table S3:** Proteins identified by mass spectrometry from hippocampal neuron intracellular proteins sample

| **Score** | **% Cov (95)** | **Uniprot id** | **Protein Name** | **Peptides (95%)** |
| --- | --- | --- | --- | --- |
| 69.65 | 42.0 | P12346 | Serotransferrin | 35 |
| 50.42 | 14.2 | Q6MG79 | Complement component 4 | 28 |
| 31.08 | 33.4 | Q5M7V3 | LOC367586 protein | 19 |
| 38.24 | 16.6 | G3V9R2 | Protein Cfh | 18 |
| 34.99 | 35.7 | P20059 | Hemopexin | 18 |
| 39.01 | 17.1 | Q91YB6 | Complement inhibitory factor H | 18 |
| 27.03 | 29.9 | Q5M842 | IgG-2a protein | 17 |
| 22.07 | 7.0 | M0RBJ7 | Complement C3 | 14 |
| 23.97 | 12.0 | P06687 | Sodium/potassium-transporting ATPase subunit alpha-3 | 11 |
| 15.80 | 16.1 | Q5I0J0 | Immunoglobulin heavy chain | 11 |
| 19.45 | 24.3 | P85108 | Tubulin beta-2A chain | 10 |
| 17.82 | 62.7 | Q05175 | Brain acid soluble protein 1 | 9 |
| 14.22 | 24.4 | Q4KM66 | LOC500183 protein | 9 |
| 15.49 | 20.7 | G3V7C6 | RCG45400 | 8 |
| 14.86 | 38.0 | I7FKL4 | Myelin basic protein transcript variant 1 | 8 |
| 12.01 | 44.3 | P01836 | Ig kappa chain C region | 8 |
| 10.70 | 55.8 | P20767 | Ig lambda-2 chain C region | 7 |
| 14.00 | 19.5 | P63259 | Actin, cytoplasmic 2 | 7 |
| 10.96 | 12.7 | F1LP05 | ATP synthase subunit alpha | 6 |
| 10.00 | 11.2 | I1T7F1 | GLT1a splice variant | 5 |
| 9.54 | 15.2 | M9MMN0 | Protein Ighg3 | 5 |
| 10.00 | 34.0 | P02091 | Hemoglobin subunit beta-1 | 5 |
| 9.67 | 10.1 | Q569B3 | Igh-6 protein | 5 |
| 7.17 | 7.2 | Q5M891 | C4b-binding protein alpha chain | 5 |
| 7.94 | 33.1 | B1H216 | Hemoglobin alpha | 4 |
| 8.01 | 14.1 | M0R660 | Protein RGD1565368 | 4 |
| 6.07 | 31.4 | P04638 | Apolipoprotein A-II | 4 |
| 8.00 | 9.5 | P09951 | Synapsin-1 | 4 |
| 6.95 | 25.2 | P11517 | Hemoglobin subunit beta-2 | 4 |
| 6.85 | 9.1 | Q99PS8 | Histidine-rich glycoprotein | 4 |
| 3.99 | 9.5 | P04636 | Malate dehydrogenase | 3 |
| 6.60 | 37.1 | P06765 | Platelet factor 4 | 3 |
| 6.21 | 7.6 | P13233 | 2',3'-cyclic-nucleotide 3'-phosphodiesterase | 3 |
| 5.11 | 22.7 | P19939 | Apolipoprotein C-I | 3 |
| 7.11 | 5.1 | P47942 | Dihydropyrimidinase-related protein 2 | 3 |
| 6.16 | 11.6 | P60203 | Myelin proteolipid protein | 3 |
| 4.13 | 6.2 | Q5M7T5 | Protein Serpinc1 | 3 |
| 5.77 | 6.7 | Q6P9V9 | Tubulin alpha-1B chain | 3 |
| 4.00 | 11.4 | F1M229 | Uncharacterized protein | 2 |
| 2.85 | 4.5 | G3V6D3 | ATP synthase subunit beta | 2 |
| 4.29 | 15.9 | M0R4L7 | Histone H2B | 2 |
| 2.00 | 6.8 | M0RBP7 | Uncharacterized protein | 2 |
| 2.76 | 6.6 | P06866 | Haptoglobin | 2 |
| 4.25 | 6.8 | P15473 | Insulin-like growth factor-binding protein 3 | 2 |
| 4.66 | 8.8 | P24090 | Alpha-2-HS-glycoprotein | 2 |
| 3.15 | 5.4 | P59215 | Guanine nucleotide-binding protein subunit alpha | 2 |
| 3.17 | 4.4 | Q5M8A0 | Kng2 protein | 2 |
| 1.72 | 4.6 | Q6IRS6 | Fetub protein | 2 |
| 1.34 | 10.8 | Q99ME0 | CXC chemokine RTCK1 | 2 |
| 2.05 | 2.7 | B0BMT0 | RCG47746, isoform CRA_a | 1 |
| 2.00 | 9.4 | D3ZC54 | Uncharacterized protein | 1 |
| 0.40 | 3.3 | D3ZCA0 | Proline synthetase co-transcribed | 1 |
| 0.79 | 0.5 | D4A8Q2 | Protein Kndc1 | 1 |
| 2.00 | 10.3 | F1LYU4 | Uncharacterized protein | 1 |
| 2.00 | 12.1 | F1LZ11 | Uncharacterized protein | 1 |
| 2.00 | 13.5 | F1M0U4 | RCG21092 | 1 |
| 0.76 | 6.4 | F1M1Q7 | Uncharacterized protein | 1 |
| 0.68 | 4.7 | G3V7N9 | Complement C1q subcomponent subunit B | 1 |
| 2.17 | 0.7 | G3V7W2 | Fibronectin | 1 |
| 2.03 | 8.5 | G3V8Z5 | Uncharacterized protein | 1 |
| 2.05 | 4.8 | M0R7B4 | Protein LOC684828 | 1 |
| 0.67 | 2.7 | M0RAS8 | Elongation factor 1-alpha | 1 |
| 2.30 | 1.7 | M0RCB1 | Uncharacterized protein | 1 |
| 2.63 | 13.7 | M0RDZ5 | Uncharacterized protein | 1 |
| 1.00 | 15.0 | O09019 | Nestin | 1 |
| 0.90 | 2.1 | P00507 | Aspartate aminotransferase | 1 |
| 0.85 | 1.0 | P02770 | Serum albumin | 1 |
| 1.66 | 2.9 | P07335 | Creatine kinase B-type | 1 |
| 1.41 | 4.6 | P07340 | Sodium/potassium-transporting ATPase subunit beta-1 | 1 |
| 1.06 | 5.9 | P10888 | Cytochrome c oxidase subunit 4 isoform 1 | 1 |
| 2.01 | 1.5 | P19527 | Neurofilament light polypeptide | 1 |
| 1.51 | 3.7 | P22057 | Prostaglandin-H2 D-isomerase | 1 |
| 1.12 | 3.0 | P24594 | Insulin-like growth factor-binding protein 5 | 1 |
| 2.00 | 3.5 | P54311 | Guanine nucleotide-binding protein G(I)/G(S)/G(T) subunit beta-1 | 1 |
| 2.00 | 1.7 | P61765 | Syntaxin-binding protein 1 | 1 |
| 0.82 | 3.2 | P61983 | 14-3-3 protein gamma | 1 |
| 2.23 | 5.7 | P63102 | 14-3-3 protein zeta/delta | 1 |
| 2.00 | 3.0 | Q6AY07 | Fructose-bisphosphate aldolase | 1 |
| 2.00 | 3.4 | Q6IRH6 | Slc25a3 protein | 1 |
| 2.00 | 4.9 | Q6MFX9 | Myelin oligodendrocyte glycoprotein | 1 |
| 2.00 | 2.2 | Q6P7S6 | Clusterin | 1 |
| 2.00 | 2.9 | Q6PAH0 | Apolipoprotein E | 1 |
| 2.00 | 1.0 | Q7TMA9 | Aa1249 | 1 |
| 2.30 | 2.0 | Q91WX0 | Complement factor H-related protein | 1 |
| 2.00 | 5.5 | Q91Y34 | Platelet phospholipase A2 | 1 |
| 2.00 | 0.8 | Q920H8 | Hephaestin | 1 |
| 2.00 | 3.0 | Q99NA5 | Isocitrate dehydrogenase [NAD] subunit alpha | 1 |
| 1.89 | 1.2 | Q9ER34 | Aconitate hydratase | 1 |
| 0.59 | 5.8 | Q9Z1I0 | Shc transforming protein | 1 |

**Table S4:** Proteins identified by mass spectrometry from hippocampal neuron plasma membrane proteins sample

| **Score** | **% Cov (95)** | **Uniprot id** | **Protein Name** | **Peptides (95%)** |
| --- | --- | --- | --- | --- |
| 32.76 | 20.8 | P06687 | Sodium/potassium-transporting ATPase subunit alpha-3 | 16 |
| 28.02 | 27.2 | P12346 | Serotransferrin | 12 |
| 22.06 | 29.9 | G3V7C6 | RCG45400 | 12 |
| 22.06 | 15.3 | P06686 | Sodium/potassium-transporting ATPase subunit alpha-2 | 10 |
| 17.44 | 8.9 | Q7TP24 | Ba1-667 | 9 |
| 17.30 | 44.1 | I7FKL4 | Myelin basic protein transcript variant 1 | 10 |
| 15.04 | 21.3 | P63259 | Actin, cytoplasmic 2 | 10 |
| 14.78 | 19.3 | V9GZ85 | Protein LOC100361457 | 10 |
| 14.28 | 23.6 | P85108 | Tubulin beta-2A chain | 11 |
| 14.13 | 17.3 | Q99PS8 | Histidine-rich glycoprotein | 7 |
| 12.00 | 14.7 | P31596 | Excitatory amino acid transporter 2 | 7 |
| 11.45 | 8.2 | Q6IMF3 | Keratin, type II cytoskeletal 1 | 6 |
| 11.23 | 12.1 | P11275 | Calcium/calmodulin-dependent protein kinase type II subunit alpha | 6 |
| 10.20 | 14.4 | Q6P9V9 | Tubulin alpha-1B chain | 5 |
| 10.00 | 47.3 | Q05175 | Brain acid soluble protein 1 | 5 |
| 9.70 | 13.0 | Q5M7V3 | LOC367586 protein | 5 |
| 9.38 | 8.7 | Q6IFW6 | Keratin, type I cytoskeletal 10 | 5 |
| 8.70 | 14.8 | P13233 | 2',3'-cyclic-nucleotide 3'-phosphodiesterase | 4 |
| 8.12 | 9.6 | Q8K5B5 | Glutamate transporter GLT1b | 6 |
| 8.00 | 9.0 | I1T7F1 | GLT1a splice variant | 4 |
| 8.00 | 11.0 | Q5I0J0 | Immunoglobulin heavy chain (Gamma polypeptide) | 4 |
| 7.53 | 10.2 | P20059 | Hemopexin | 4 |
| 6.39 | 17.2 | D4ABT0 | Guanine nucleotide-binding protein G(o) subunit alpha | 3 |
| 6.09 | 6.7 | Q5XIF6 | Tubulin alpha-4A chain | 3 |
| 6.06 | 13.8 | G3V6D3 | ATP synthase subunit beta | 3 |
| 6.05 | 22.3 | P62804 | Histone H4 | 3 |
| 6.00 | 5.6 | P61765 | Syntaxin-binding protein 1 | 3 |
| 6.00 | 17.1 | Q4KM66 | LOC500183 protein | 3 |
| 5.52 | 7.0 | P09951 | Synapsin-1 | 3 |
| 5.31 | 3.0 | Q6P6Q2 | Keratin, type II cytoskeletal | 3 |
| 5.12 | 19.7 | P02091 | Hemoglobin subunit beta-1 | 3 |
| 5.03 | 34.0 | P01836 | Ig kappa chain C region, A allele | 3 |
| 4.92 | 13.5 | P47819 | Glial fibrillary acidic protein | 2 |
| 4.52 | 5.6 | B0BMT0 | RCG47746, isoform CRA_a | 2 |
| 4.44 | 15.9 | M0R4L7 | Histone H2B | 3 |
| 4.28 | 22.9 | P06765 | Platelet factor 4 | 2 |
| 4.25 | 19.8 | M0R660 | Protein RGD1565368 | 2 |
| 4.13 | 11.6 | P60203 | Myelin proteolipid protein | 3 |
| 4.11 | 28.2 | Q71DI1 | Dermcidin | 3 |
| 4.07 | 4.6 | D3ZQQ5 | Dynamin-1 | 2 |
| 4.05 | 15.8 | P07340 | Sodium/potassium-transporting ATPase subunit beta-1 | 2 |
| 4.00 | 13.5 | P54311 | Guanine nucleotide-binding protein G(I)/G(S)/G(T) subunit beta-1 | 3 |
| 4.00 | 13.7 | G3V7Y3 | ATP synthase subunit delta | 2 |
| 4.00 | 21.1 | M0RDZ5 | Uncharacterized protein | 2 |
| 4.00 | 8.3 | P60881 | Synaptosomal-associated protein 25 | 2 |
| 4.00 | 22.6 | Q63754 | Beta-synuclein | 2 |
| 4.00 | 9.4 | Q6P9Y4 | ADP/ATP translocase 1 | 2 |
| 4.00 | 5.0 | Q812E9 | Neuronal membrane glycoprotein M6-a | 2 |
| 3.74 | 9.9 | P01830 | Thy-1 membrane glycoprotein | 2 |
| 3.22 | 15.4 | Q6I8Q6 | Histone H2A | 2 |
| 3.11 | 17.1 | P63102 | 14-3-3 protein zeta/delta | 2 |
| 2.80 | 4.0 | M0RBJ7 | Complement C3 | 1 |
| 2.52 | 6.5 | G3V8Q2 | Alpha-internexin | 1 |
| 2.41 | 17.8 | Q9WUW2 | Vesicle associated membrane protein 2B | 1 |
| 2.36 | 14.8 | R9PY00 | Vesicle-associated membrane protein 2 | 1 |
| 2.27 | 8.1 | P00762 | Anionic trypsin-1 | 6 |
| 2.27 | 1.0 | Q5EBC0 | Inter alpha-trypsin inhibitor, heavy chain 4 | 1 |
| 2.22 | 34.3 | P37377 | Alpha-synuclein | 1 |
| 2.20 | 1.6 | Q6IFU8 | Keratin, type I cytoskeletal 17 | 1 |
| 2.11 | 17.2 | P10888 | Cytochrome c oxidase subunit 4 isoform 1 | 1 |
| 2.04 | 6.1 | P19527 | Neurofilament light polypeptide | 1 |
| 2.04 | 4.2 | P47942 | Dihydropyrimidinase-related protein 2 | 1 |
| 2.04 | 8.1 | Q6IRH6 | Slc25a3 protein | 1 |
| 2.00 | 12.4 | P12075 | Cytochrome c oxidase subunit 5B | 2 |
| 2.00 | 22.0 | B2RZD6 | Ndufa4 protein | 1 |
| 2.00 | 0.4 | D3ZZ51 | Protein Prrc2b | 1 |
| 2.00 | 10.2 | M0R7G5 | Protein LOC100911905 | 1 |
| 2.00 | 2.7 | M0RAS8 | Elongation factor 1-alpha | 1 |
| 2.00 | 15.0 | O09019 | Nestin | 1 |
| 2.00 | 5.3 | P06349 | Histone H1t | 1 |
| 2.00 | 4.7 | P20788 | Cytochrome b-c1 complex subunit Rieske | 1 |
| 2.00 | 16.9 | P29419 | ATP synthase subunit e | 1 |
| 2.00 | 2.4 | P62632 | Elongation factor 1-alpha 2 | 1 |
| 2.00 | 4.9 | Q5RK07 | Igh-6 protein | 1 |
| 2.00 | 4.7 | Q6AY07 | Fructose-bisphosphate aldolase | 1 |
| 2.00 | 3.1 | S5RZM8 | Cytochrome c oxidase subunit 2 | 1 |
| 1.89 | 3.8 | P11980 | Pyruvate kinase isozymes M1/M2 | 1 |
| 1.70 | 1.4 | P62815 | V-type proton ATPase subunit B | 1 |
| 1.66 | 3.6 | P63012 | Ras-related protein Rab-3A | 1 |
| 1.64 | 3.3 | P07825 | Synaptophysin | 1 |
| 1.55 | 3.8 | F1M882 | Secretory carrier-associated membrane protein 5 | 1 |
| 1.46 | 1.0 | F1MA98 | Nucleoprotein TPR | 1 |
| 1.42 | 1.6 | F1LPY7 | Trans-2-enoyl-CoA reductase, mitochondrial | 1 |
| 1.22 | 6.3 | B1H216 | Hemoglobin alpha, adult chain 2 | 1 |
| 1.16 | 0.0 | Q9QUL6 | Vesicle-fusing ATPase | 1 |
| 1.07 | 8.9 | P11240 | Cytochrome c oxidase subunit 5A | 1 |
| 0.77 | 5.6 | Q0QF43 | Malate dehydrogenase | 1 |
| 0.75 | 2.6 | D4A4M0 | Protein Shisa6 | 1 |
| 0.71 | 0.7 | D3ZJF8 | Protein Fcgbp | 1 |
| 0.54 | 0.5 | P21571 | ATP synthase-coupling factor 6, mitochondrial | 1 |
| 0.51 | 0.5 | P19527 | Neurofilament light polypeptide | 1 |
| 0.50 | 0.5 | P61983 | 14-3-3 protein gamma | 1 |
| 0.44 | 0.5 | P21707 | Synaptotagmin-1 | 1 |
| 0.36 | 0.5 | D3ZKH2 | Protein Mon2 | 1 |
| 0.31 | 0.5 | D3Z9Z2 | Putative lipoyltransferase 2, mitochondrial | 1 |
| 0.29 | 0.5 | Q9Z1I0 | Shc transforming protein | 1 |
| 0.23 | 0.5 | P20767 | Ig lambda-2 chain C region | 1 |
| 0.21 | 0.5 | F1LXD6 | Protein Pitpnm2 | 1 |
| 0.20 | 0.5 | Q66HG1 | Cyclin-D-binding Myb-like transcription factor 1 | 1 |
| 0.10 | 0.5 | G3V733 | Synapsin-2 | 1 |
| 0.09 | 0.5 | D4ABV5 | Calmodulin | 1 |
| 0.08 | 0.5 | D3ZV39 | Protein Armcx4 | 1 |
| 0.08 | 0.5 | M0RBX6 | Histone H3 | 1 |
| 0.08 | 0.5 | R9PXT6 | Focal adhesion kinase 1 | 1 |
| 0.07 | 0.5 | D4AA54 | Protein Plekhh1 | 1 |
| 0.06 | 0.5 | Q6PAH0 | Apolipoprotein E | 1 |
| 0.05 | 0.5 | D3ZSV6 | DEAH (Asp-Glu-Ala-His) box polypeptide 33 (Predicted) | 1 |
| 0.05 | 0.5 | D4A3W6 | Dual specificity phosphatase 16 (Predicted) | 1 |
| 0.05 | 0.5 | D4A9W1 | Protein Ccdc88c | 1 |
| 0.05 | 0.5 | G3V984 | Protein bassoon | 1 |
| 0.05 | 0.5 | Q71S46 | ATP synthase F(0) complex subunit C3, mitochondrial | 1 |
